# Complexity matching: brain signals mirror environment information patterns during music listening and reward

**DOI:** 10.1101/693531

**Authors:** Sarah M. Carpentier, Andrea R. McCulloch, Tanya M. Brown, Petra Ritter, Zhang Wang, Valorie Salimpoor, Kelly Shen, Anthony Randal McIntosh

## Abstract

Understanding how the human brain integrates information from the environment with ongoing, internal brain signals in order to produce individual perspective is an essential element of understanding the human mind. Brain signal complexity, measured with multiscale entropy, has been employed as a measure of information processing in the brain (Carpentier et al., 2016), and we propose that it can also be used to measure the information available from a stimulus. We can directly assess the correspondence, or functional isomorphism, between brain signal complexity and stimulus complexity as an indication of how well the brain reflects the content of the environment in an analysis that we termed *complexity matching*. Music makes an ideal stimulus input because it is a multidimensional, complex signal, and because of its emotion and reward-inducing potential. We found that electroencephalography (EEG) complexity was lower and more closely resembled the musical complexity when participants performed a perceptual task that required them to closely track the acoustics, compared to an emotional task that asked them to think about how the music made them feel. Music-derived reward scores on the Barcelona Music Reward Questionnaire (Mas-Herrero et al., 2013) correlated with worse complexity matching and higher EEG complexity. Compared to perceptual-level processing, emotional and reward responses are associated with additional internal information processes above and beyond those in the external stimulus.

**Significance Statement:** Experience of our world is combination of the input from the environment, our expectations, and individual responses. For example, the same piece of music can elict happiness in one person and sadness in another. We researched this by measuring the information in pieces of music and whether listener’s brain more closely followed that, or whether additional information was added by the brain. We noted when listener’s were reacting to how music made them feel, their brains added more information and the degree to which this occurred related to how much they find music rewarding. Thus, we were able to provide clues as to how the brain integrates incoming information, adding to it to provide a richer perceptual and emotional experience.

---

Some contemporary theories suggest that functional brain networks engage and disengage to integrate information during cognitive processes (Tononi et al., 1994; McIntosh, 2000; Bressler and Kelso, 2001). This network activity generates highly variable and complex brain signals; therefore, brain signal complexity can serve as an indicator of the information processing of the system (Deco, Jirsa, & McIntosh, 2011; Ghosh et al., 2008; McIntosh, Kovačević, & Itier, 2008). Accordingly, brain signal complexity is higher during states of greater knowledge representation (e.g. Heisz, Shedden, & McIntosh, 2012), and increased following longitudinal music training (Carpentier et al., 2016). The present study measured signal complexity to investigate whether there is a correspondence between information patterns in brain signals and those in the individual’s environment. Furthermore, we were interested in whether this correspondence would be related to the cognitive-affective state of the individual.

Music offers an ideal stimulus from which to measure information content. Complicated music structures are created following application of combination rules to subordinate motifs. This makes it possible to describe a functional isomorphism between the brain and environment information by calculating music complexity and comparing it to the complexity of brain signals of the listener. We propose that this measure of comparison between EEG complexity and music complexity, or *complexity matching*, will provide a relative indication of the degree to which environmental information structure is reflected in brain signal structure.

Complexity matching is calculated as Procrustes distance (Gower, 1975) between the music and EEG multiscale entropy (MSE). MSE calculates sample entropy at multiple timescales (Costa et al., 2002, 2005). Like brain signals, music also has structure at multiple timescales, and MSE seems an appropriate measure of complexity for a given music passage. Procrustes distance produces a quantity of similarity between the structure of the music and the structure of the ensuing brain signals. A relatively high value of matching together with lower EEG MSE would suggest that the brain has activated the necessary information processing resources for immediate perception, and little else. Conversely, relatively lower complexity matching but high neural complexity would suggest that internal processes, different from immediate stimulus perception, dominate the neural response. This metric was inspired by the ideas in Tononi et al. (Tononi et al., 1996)

The level of brain-environment information integration may be related to cognition and subjective perspective. To examine the relationship between brain-environment matching and cognitive-affective state, we calculated complexity matching while participants performed a music perception task and a music emotion evocation task. We expect that active attention to the acoustics of the music during the perceptual task will be accompanied by brain signal complexity that more closely resembles the music, compared to the emotional task that involves additional internally processes and, therefore, will provide less of a match to the environment.

We also analyzed the relationship between complexity matching and music-derived reward scores from the Barcelona Music Reward Questionnaire (BMRQ). The ability to perceive musical structure is essential to the enjoyment of music (Meyer, 1956; Huron, 2006): therefore, it is possible that a certain minimum quantity of complexity matching may be required for the listener to have the necessary appreciation of the underlying ‘gist’ or skeleton structure of the piece. Perhaps without sufficient neural integration of music signals, the listener would be unable to perceive separate noise sounds as unified. In accordance with our prediction that the emotional task will be associated with a lower complexity match than the perceptual task, we expect that higher music reward involves internally-driven, individual responses and therefore will correlate with higher complexity and lower complexity matching.

## Materials and Methods

### Participants

Eighteen healthy young adults aged 19-35 (M = 26; 10 female) were recruited from the Greater Toronto Area to take part in the study and provided written informed consent in accordance with the joint Baycrest Centre-University of Toronto Research Ethics Committee. Prior to arriving to the lab for the experimental session, participants completed an online questionnaire about their music listening habits, and musical training was assessed as a 1-5 scale: 1) No formal training, cannot play an instrument; 2) Can play an instrument without formal training; 3) Less than 1 year of formal music training; 4) Between 1-5 years of formal training; and 5) More than 5 years of formal training.

### Barcelona Music Reward Questionnaire

Music reward is highly individual, and the BMRQ was developed to describe some of the main facets of the variance in how people experience reward from music listening (Mas-Herrero et al., 2013). Participants are asked to indicate the level of agreement with each of 20 statements by using a 5-point scale ranging from (1) “fully disagree” to (5) “fully agree,” with a higher score indicating the subject experiences more music reward and a lower score indicating they do not experience music associated rewarding feelings. These statements represent five major factors of music reward: (1) Emotional Evocation; (2) Mood Regulation; (3) Musical Seeking; (4) Social Reward; and (5) Sensory-Motor.

*Emotional Evocation* refers to the idea that music can both convey and induce emotion (also referred to as emotional contagion), such as joy or sadness, and that listeners might seek out music that contains emotion (Juslin and Laukka, 2004; Juslin and Västfjäll, 2008; Vuoskoski and Eerola, 2012). The BMRQ distinguishes evoked feelings, which may be short-lived and vary across a single music piece; from the way some listeners use music to alter their own longer lasting mood or hedonic state after the song has finished (e.g. Carter, Wilson, Lawson, & Bulik, 1995; Västfjäll, 2001). *Mood Regulation* refers to the idea that music can be used to comfort, relieve stress, or enhance relaxation (for a review see Juslin & Sloboda, 2010), and a particular point has been raised about the use of music in marketing or film to manipulate and induce hedonic states (Cohen, 2001). *Musical Seeking* can also be referred to as “knowing about music.” This facet describes that some listeners get reward from extracting, pursuing, sharing, and seeking information regarding specific music pieces, composers, performers, or other information related to music. Listeners may also experience pleasure when recognizing music quotations or allusions to other works. *Social Reward* may be gained by music through its enhancement of social bonds or social cohesion (Cross and Morley, 2009). Lastly, the *Sensory-Motor* facet captures reward experienced by the pull music has over some people to move to music.

### Behaviour Tasks

Forty operatic and classical musical segments were selected after piloting for a range of emotional reactions of the listener and of pitch and tempo. The pieces spanned a range with only instruments to both instruments and voice. For this study, we wanted to ensure the range of individual experience was as broad as possible to get reasonable ranges of arousal and valence ratings and comparable volatility in the perceptual task. Segment lengths ranged between 0:40-1:17 min. This choice was made to allow each segment to conclude naturally at the end of a musical phrase, rather than ending abruptly in the middle. Thirty pieces were selected for the emotional task and ten pieces for the perceptual task (Table 1). Importantly, there was no difference in the music complexity (MSE) between the tasks (*p* > 0.1).

**Table 1.**
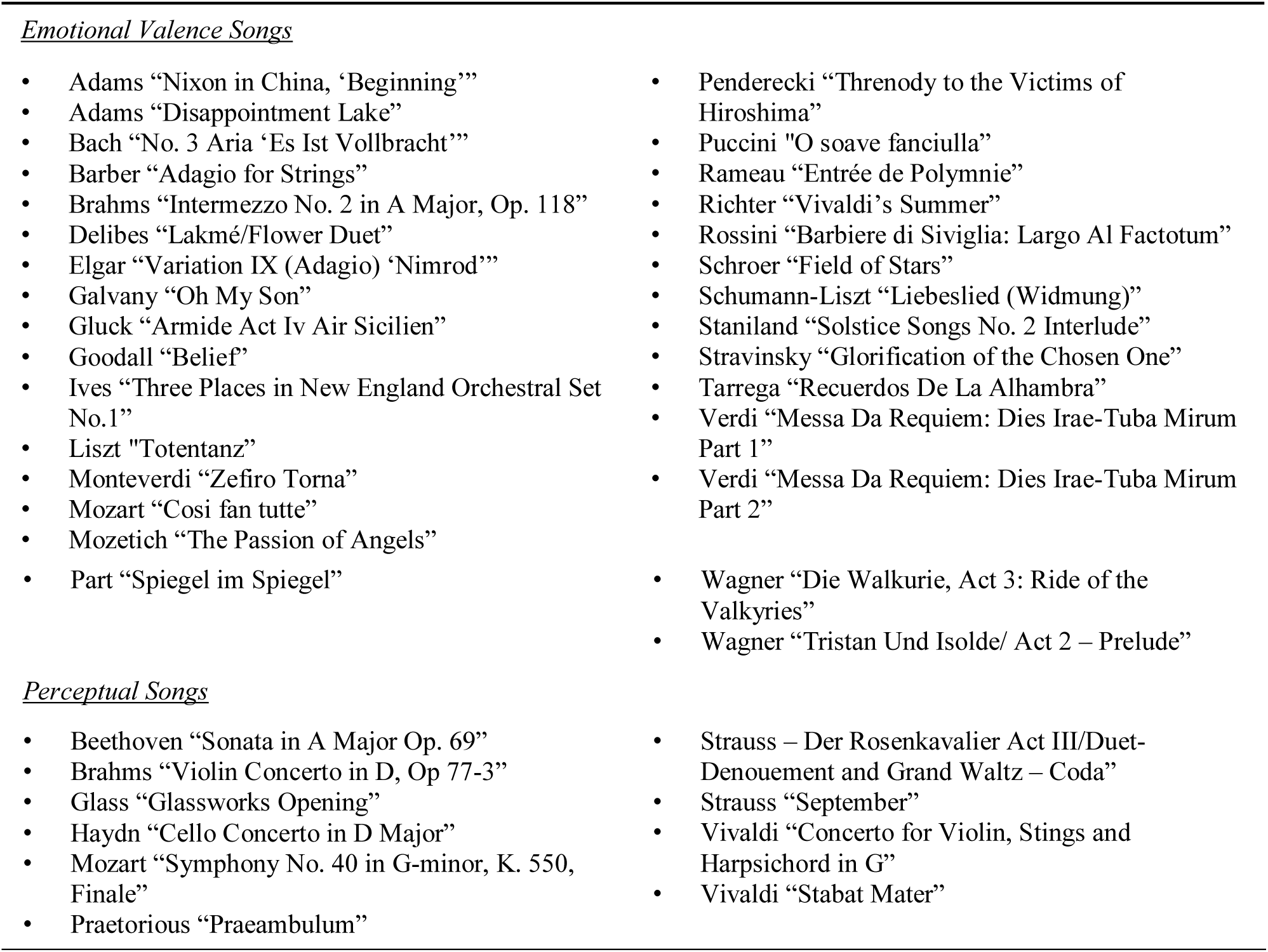
List of songs for each tasks.

During the emotional task, a computer screen in front of participants showed four quadrants marked on two dimensions: Stimulating-Relaxing, and Pleasant-Unpleasant (Figure 1). Participants were asked to move a mouse around the quadrant space in a continuous manner during each song based on *how the music made them feel* on the two dimensions. Participants were explicitly instructed to report of their own feelings during music listening, and not the alternative of reporting on what emotions they believe are expressed in the music (emotional conveyance). The task design was modeled after the *valence-arousal model* of Hunter & Schellenberg, 2010. They labeled their dimensions high arousal-low arousal and positive valence-negative valence, and we altered our labels after pilot tests to be more intuitive for subjects. This valence-arousal model is designed to capture a wide range of emotions. In their study, difference valence and arousal combinations were associated with multiple different emotions. For example, high arousal/negative valence was correlated with distress, fear and anger, low arousal/positive valence was associated with feelings of peace, contentment and relaxation. Participants from our pilot sample gave similar reports. In this way, it is possible to capture a larger range of emotions without limiting responses to more specific emotions.

**Figure 1.**
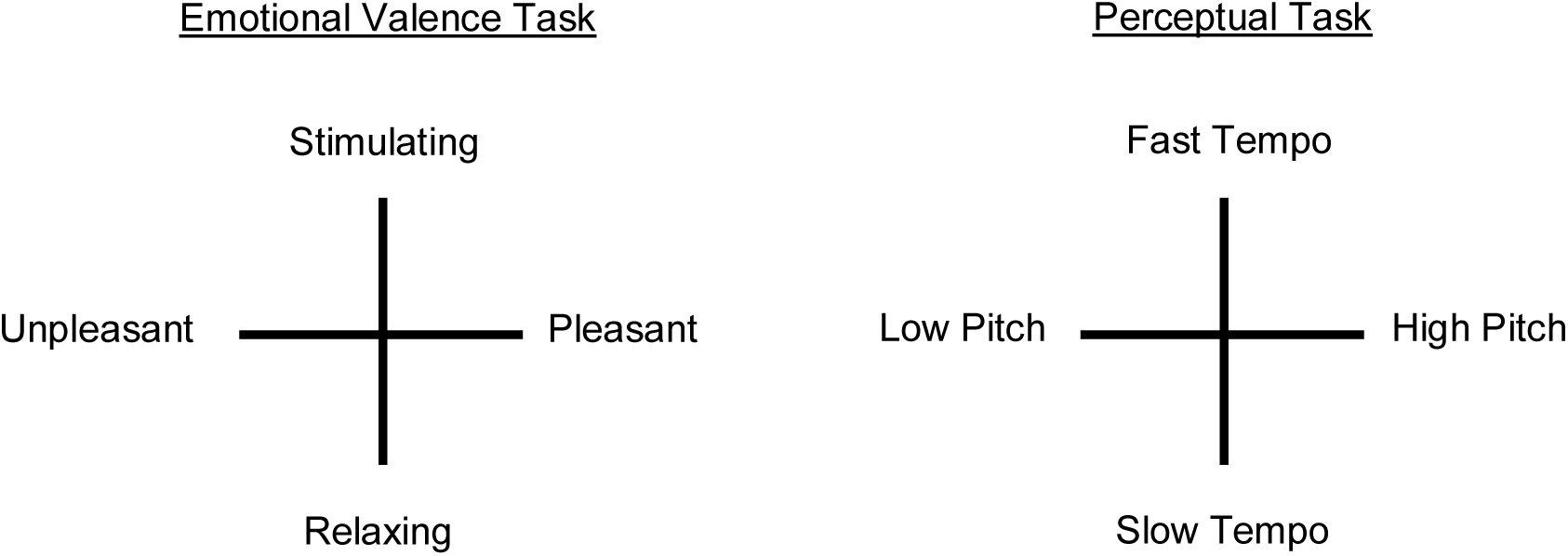
Participants viewed screens with each of the above quadrants during each task. They were asked to move a mouse continuously around the quadrant space depending on how the music was making them feel in that moment (emotional task) or based on their judgments of pitch and tempo (perceptual task).

The perceptual task mimicked the emotional task, with the difference being participants were required to assess pitch and tempo for each song (Figure 1). Once again, a screen in front of them displayed four quadrants with two dimensions (High-Low Pitch and Fast-Slow Tempo), and participants moved a mouse in a continuous manner on the screen based on the pitch and tempo of each song.

### Experimental Procedure

The experimental session began with five perceptual task songs, followed by all thirty emotional songs, and concluded with the remaining five perceptual songs. Perceptual songs were always presented in the same order. Emotional songs were presented in one of two counterbalance orders. Pieces in the first order were curated to have a sense of flow between them and avoid jarring transitions from one song to the next that may disrupt emotional experiences. The second order was the reverse of the first. There was no significant effect of counterbalance order on any of our measures. All stimuli were presented through ER 3A insert earphones (Etymotic Research, Elk Grove, U.S.A.), while participants were seated in a soundproof room.

#### EEG Recording and Pre-Processing

EEG was recorded using a 64+10 Biosemi Active Two System at a sampling rate of 512 Hz. Continuous EEG recordings were bandpass filtered at 0.5-90 Hz, with a notch filter at 55-65 Hz for line noise. The shortest music segment was 40 seconds, so EEG data for each song was segmented into 4 × 10 s epochs and baseline corrected based on a 200 ms pre-stimulus interval. Trials with excessive signal amplitude were rejected. Ocular and muscle artifact removal was performed on the remaining concatenated trials using Independent Component Analysis (ICA) implemented in EEGLAB (Delorme and Makeig, 2004). The highest number of trials lost for any subject was 8 out of 40, 7 subjects retained all trials, and the average number rejected trials from remaining subjects was 2.67, with no difference in trial rejection between conditions.

We performed source estimation at the 68 ROIs of the Desikan-Killiany Atlas (Desikan et al., 2006), using sLORETTA (Pascual-Marqui, 2002) as implemented in Brainstorm (Tadel et al., 2011). Brainstorm is documented and freely available for download under the GNU general public license (http://neuroimage.usc.edu/brainstorm). Source reconstruction was constrained to the cortical mantle of the brain template MNI/Colin27 defined by the Montreal Neurological Institute (Holmes et al., 1998). Current density for one source orientation (X component) was estimated for 15,768 equally spaced vertices and the source waveform was mapped at the 68 brain regions of interest as an average taken over all vertices in each region. Multiscale Entropy was calculated on the source waveform at each ROI for each subject as a measure of brain signal complexity.

### Data Analyses

#### Multiscale Entropy

MSE has been previously validated as a measure of brain signal complexity (Catarino et al, 2011; McIntosh et al., 2008; Mišić, Mills, Taylor, & McIntosh, 2010). We calculated MSE in two steps using the algorithm available at www.physionet.org/physiotools/mse. First, the source EEG and music signals were progressively down-sampled into multiple coarse-grained timescales where, for scale *t*, the time series is constructed by averaging the data points with non-overlapping windows of length *t*. Each element of the coarse-grained time series, 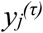, is calculated according to Eq. (2):

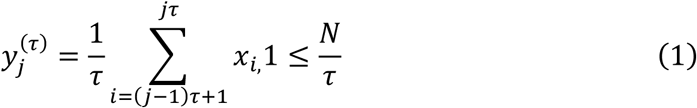

The number of scales is determined by a function of the number of data points in the signal and MSE was calculated for 100 timescales [sampling rate (512Hz) * epoch (10,000 ms)/50 time points per epoch = maximum of 102.4 scales].

Second, the algorithm calculates the sample entropy (S_E_) for each coarse-grained timeseries 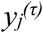:

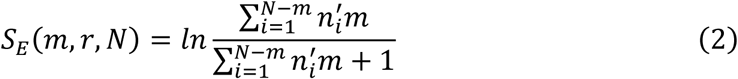

Sample entropy quantifies the predictability of a time series by calculating the conditional probability that any two sequences of *m* consecutive data points that are similar to each other within a certain similarity criterion (*r*) will remain similar at the next point (*m*+*1*) in the data set (N), where N is the length of the time series (Richman & Moorman, 2000). In this study, MSE was calculated with pattern length set to m = 2, and similarity criterion was to r = 0.5. The value r is defined as a proportion of the standard deviation of the original data (Costa, Goldberger, & Peng, 2004; Richman & Moorman, 2000). MSE estimates were obtained for each participant’s EEG source time series as a mean across single-trial entropy measures for each timescale.

Music pieces were imported into Matlab using the *wavread* function at a sampling rate of 11.25 kHz (MathWorks, Inc. Release 2011b). Music auditory signal MSE was subsequently calculated with the same parameter values and the same number of timescales as the EEG source MSE.

#### Complexity Matching

Complexity matching applies Procrustes analysis to measure the equivalence of the MSE curve for the auditory signal of a song (*X*_*1*_) and the MSE curve of the EEG source time series of a participant listening to that song (*X*_*2j*_), for all *j* ROIs individually (Gower, 1975). It minimizes the sum of the squared deviations between matching corresponding points (landmarks) from each of the two data sets (MSE curves), allowing for scaling, translation and orthogonal rotation of *X*_*1*_ to fit *X*_*2j*_, where choice of label *X*_*1*_ or *X*_*2*_ is arbitrary. *X*_*1*_ and *X*_*2j*_ must have the same number of *i* sample points, or ‘landmarks’, and Procrustes matches *X*_*1i*_ to *X*_*2ij*_. In our simple case of two vectors, the rotation matrix *T* such that *X*_*1*_ best fits *X*_*2j*_ is given as *T = V’U* from the singular value decomposition *X*_*1*_’ *X*_*2j*_ = *U’SV*. Without translation and scaling this problem is known as Procrustes rotation. Dissimilarity of *X*_*1*_ and *X*_*2j*_ is given as the *Procrustes distance*:

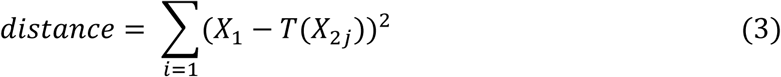

A smaller distance value denotes greater similarity between the two curves, or a closer match between them. The analysis returns a distance value for each ROI for each participant. Procrustes distance was calculated using the Matlab function *procrustes* (MathWorks, Inc. Release 2011b).

Figure 2 presents a conceptual depiction of our implementation of *complexity matching.*

**Figure 2.**
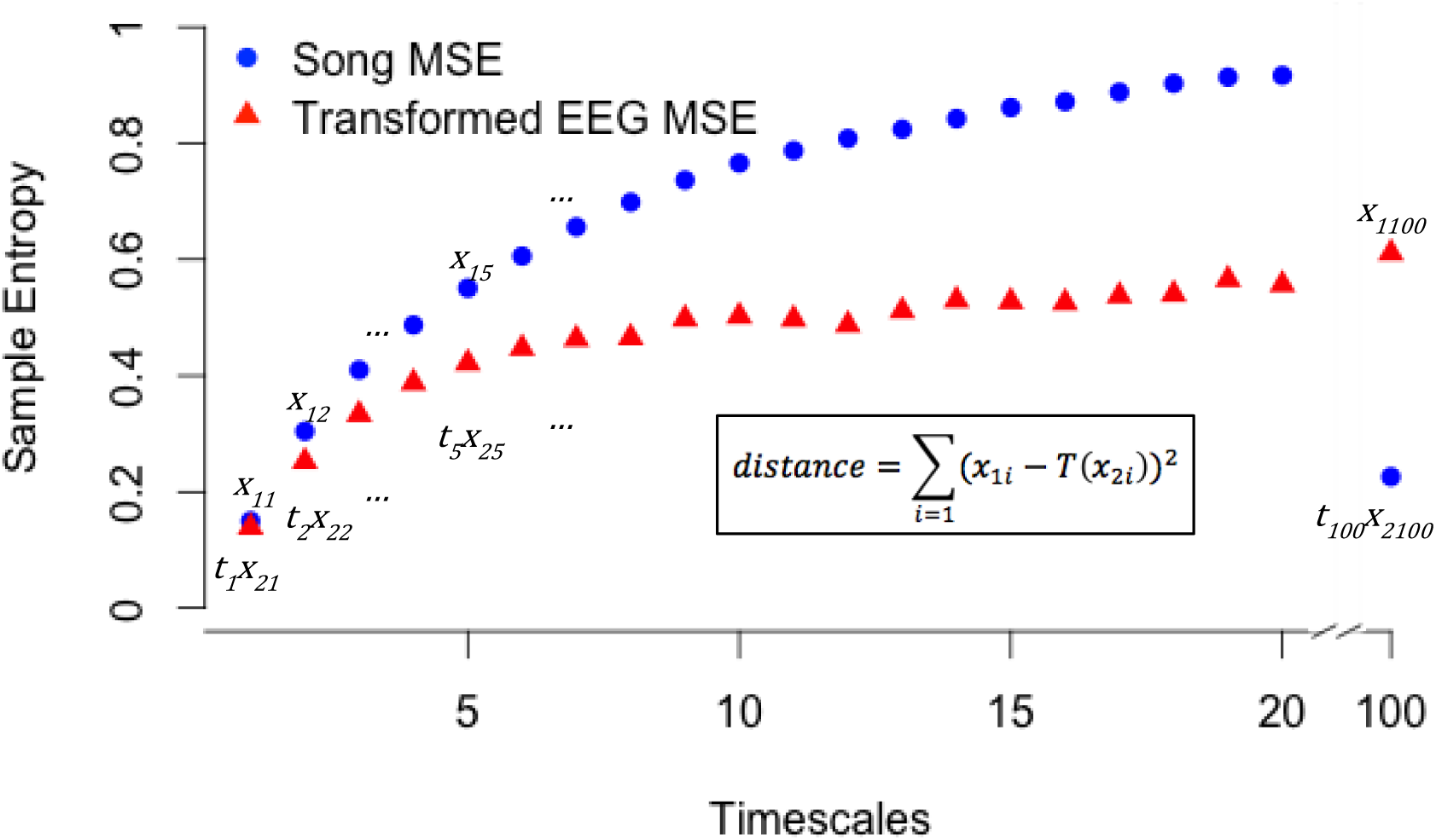
Complexity matching applies Procrustes’ analysis to determine a linear transformation (scaling, translation and orthogonal rotation) of the points in X_2_ (EEG source MSE for a given ROI) to best match the points in X_1_ (song MSE). The goodness-of-fit criterion is the sum of squared errors, and Procrustes distance is the minimized value of this dissimilarity measure. Distance is standardized by a measure of the scale of X_1_.

#### Spectral Power

Studies have found that MSE and power spectrum density (PSD) provide complementary information on neural signals (Gudmundsson et al., 2007; McIntosh et al., 2008; Mišić et al., 2010). For example, both measures follow similarities for time maturational changes, but with different spatial and temporal patterns (McIntosh et al., 2008; Lippé et al., 2009; Mišić et al., 2010). Mišić and colleagues (2014) found substantial differences between PSD and MSE effects. In their sample, individuals with Autism Spectrum Disorder (ASD) displayed only group main effects on PSD, but a group x task interaction on MSE, and the effects were different both spatially and temporally. This indicates that MSE captures an aspect of neural information processing in ASD above and beyond what can be gleaned from a traditional analysis of spectral power.

To determine the extent to which training- and task-based differences in MSE are related to spectral density, we computed PSD for all single-trial time series. Single-trial power spectra were computed using the Fast Fourier Transform. To capture the relative contribution from each frequency band, all time series were first normalized to mean = 0 and SD = 1. Given the sampling rate of 512 Hz and 5,120 data points per trial, the frequency resolution was effectively 0.100 Hz and the analysis was constrained to the [0.100, 90] Hz range, with a notch filter for line noise at 55-65 Hz.

#### Partial Least Squares

Task partial least squares analysis (PLS) was used to statistically assess task and epoch related effects in MSE and PSD. Task PLS is a multivariate statistical technique similar to canonical correlation which employs singular value decomposition (SVD) to extract latent variables (LVs) that capture the maximum covariance between the task design and neural activity. Each LV consisted of: (1) a singular vector of design scores, (2) a singular vector of saliences showing the distribution across brain regions and sampling scales, (3) a singular value (*s*) representing the covariance between the design scores and the singular image (McIntosh et al., 1996; McIntosh and Lobaugh, 2004).

The statistical significance of each LV was determined using permutation testing (Good, 2000; McIntosh and Lobaugh, 2004). The rows of *X* are randomly reordered (permuted) and the new data were subjected to SVD as before, to obtain a new set of singular values. This procedure was repeated 500 times to generate a sampling distribution of singular values under the null hypothesis that there is no association between neural activity and the task. An LV was considered significant if a singular value equal to or greater than that of the LV was present less than 5% of the time in random permutations (i.e. *p* < 0.05).

The reliability of each statistical effect was assessed through bootstrap estimation of standard error confidence intervals of the singular vector weights in each LV (Efron and Tibshirani, 1986). Random sampling with replacement of participants within conditions generated 500 bootstrap samples. In the present study, this process allowed for the assessment of the relative contribution of brain regions and timescales to each LV. Brain regions with a salience weight over standard error ratio > 3.0 correspond to a 99% confidence interval and were considered to be reliable (Sampson et al., 1989).

Finally, the dot product of an individual subject’s raw MSE data and the singular image from the LV produces a brain score. The brain score is similar to a factor score that indicates how strongly an individual subject expresses the patterns on the latent variable and allowed us to estimate 95% confidence intervals for the effects in each group and task condition.

Behavioural PLS (bPLS) is a variation on task PLS for analyzing the relationship between brain measures and the behaviour (McIntosh and Lobaugh, 2004; Krishnan et al., 2011). Similar to task PLS the SVD results in mutually orthogonal LVs, where each LV contains 1) a singular vector of saliences for the behavioural measures, (2) a singular vector of saliences for brain activity, (3) a singular value (*s*) representing the covariance between the behaviour scores and the singular image. Behaviour saliences indicate task-dependent differences and brain saliences indicate ROI-dependent differences in the brain-behaviour correlation.

## Results

One participant was excluded from the study for mild hearing loss determined by audiogram, and one other was removed for excessive motion during EEG recording, leaving N=16 subjects. Two participants did not complete the music-training questionnaire, leaving 14 subjects. For music training, of the N=14 participants: 7 reported no formal training, 1 reported can play an instrument without formal training, 1 reported less than 1 year of formal music training, 2 reported between 1-5 years of formal training, and 4 reported more than 5 years of formal training. Thus, this sample does not include a sufficient number of participants who fulfill the common requirements for musicianship (e.g. at least 10 years of formal music training, Fujioka et al., 2004), and we did not proceed with analysis of the effects of music training.

Multiscale entropy curves of sound signals from a sample of the songs are visualized in Figure 3 for illustration purposes.

**Figure 3.**
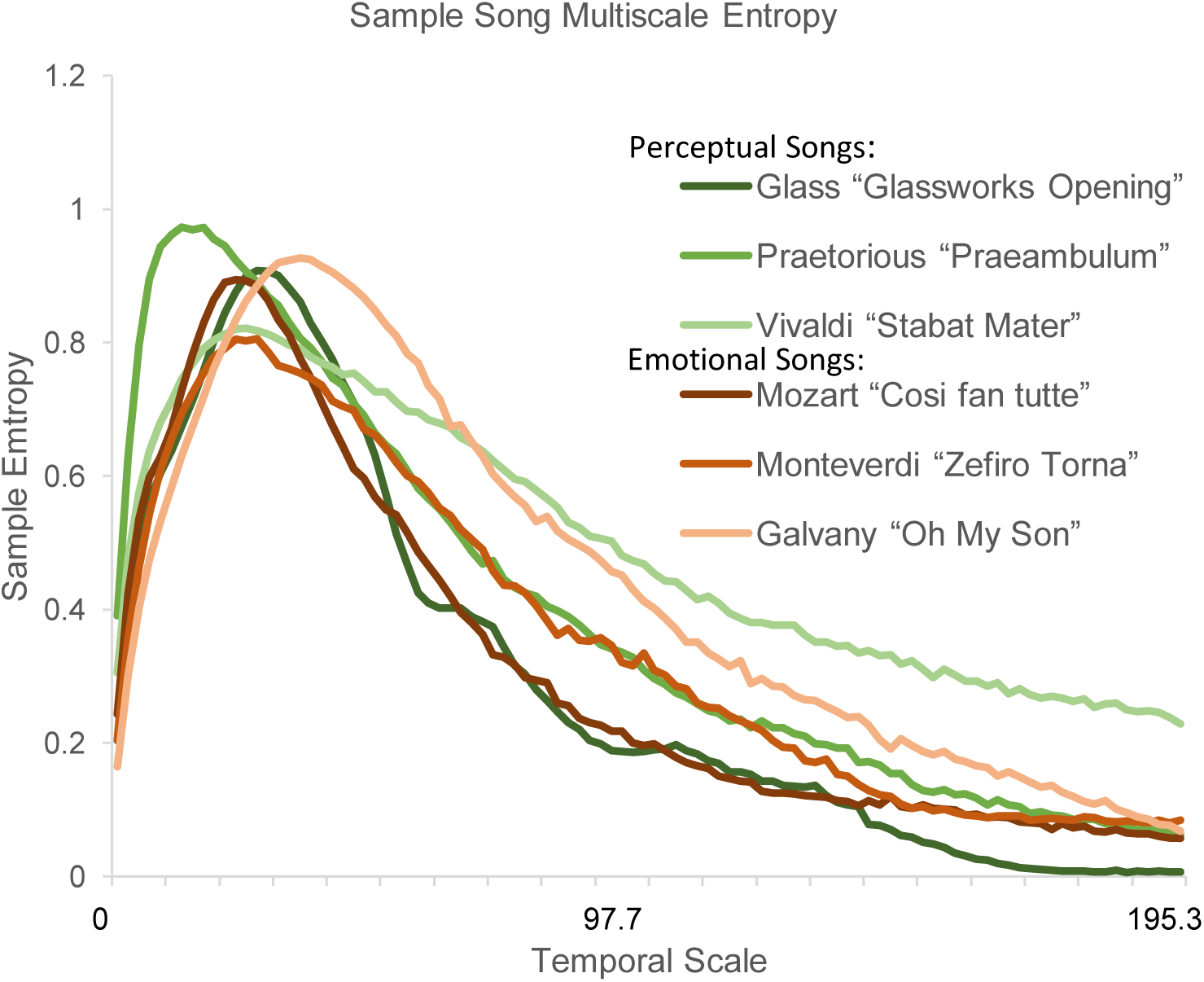
Exemplary multiscale entropy values obtained from the sound signals of a small sample of the songs from each group. Temporal scale, in milliseconds, refers to the number of data points averaged within non-overlapping windows, hence the left most values represent fine temporal scales and right more coarse scales.

### Emotional and Perceptual Tasks

We did not observe any within-task effects of emotional (e.g. stimulating compared to relaxing) or pitch/tempo (fast compared to slow) dimension ratings on any of our brain measures (MSE, Procrustes distance or PSD; all PLS *p* > .10). This may be due to the high level of variance between subjects’ emotional responses (Figure 4), or because the continuous nature of the behaviour ratings is not well suited to the dichotomization necessary for the present types of analyses.

**Figure 4.**
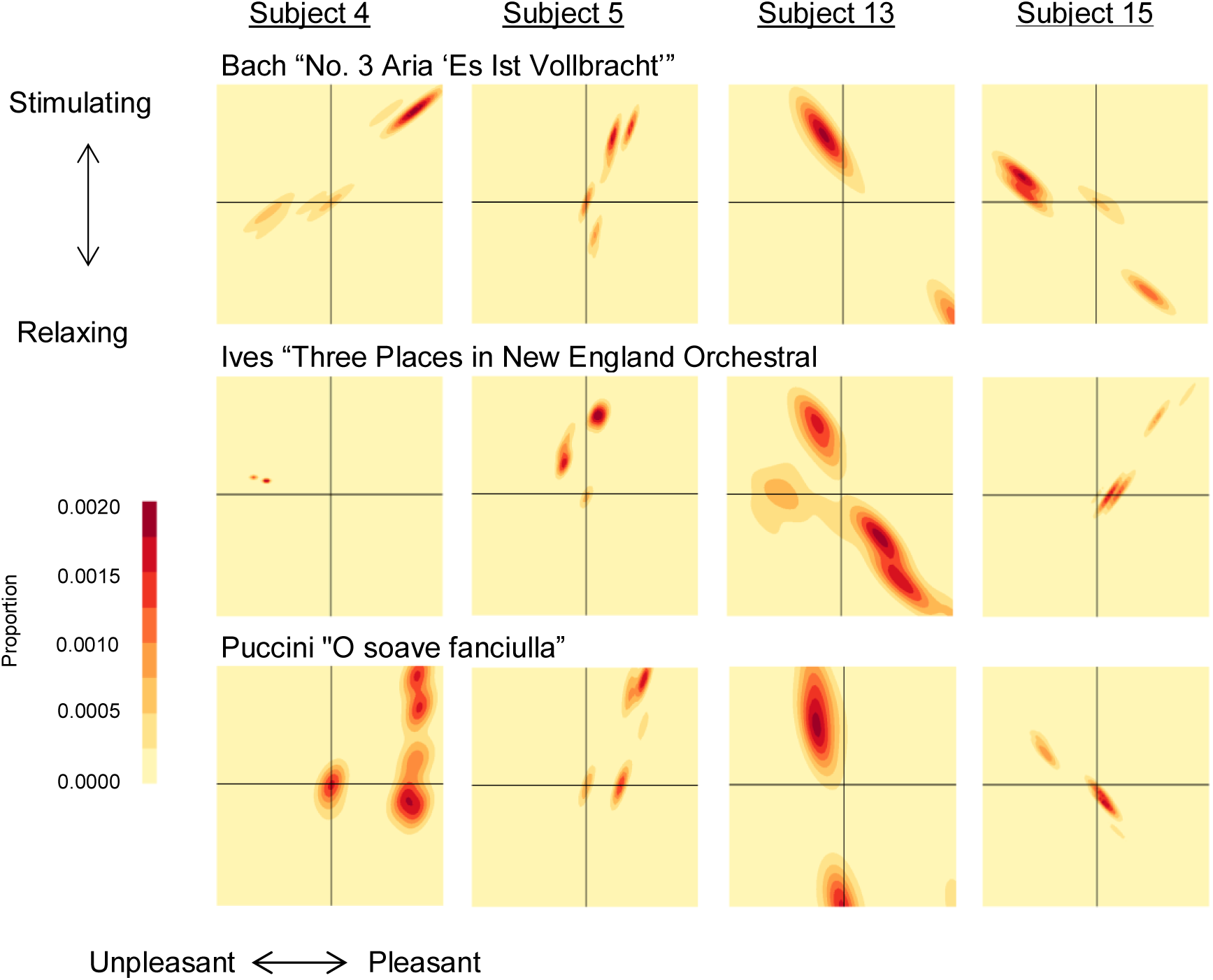
Example single-subject, single-song behaviour heat maps for the emotional valence task. Note the variation between individuals in both the valence felt and the variability of the valence within each participant and song (i.e. some participants were stable in one quadrant while some participants felt a greater range during the same song).

Examining both tasks across all four epochs, the emotional task was generally associated with higher EEG source MSE at time scales below 20, compared to the perceptual task that showed higher MSE at coarser timescales (>40) (LV = 1, p << 0, Singular Value = 1.81, 31.6% cross-block covariance; Figure 5; Figure 6). Both tasks showed an increase in MSE at finer timescales (<20) and a decrease in coarse scale MSE across epochs from the beginning to the end of the piece of music. The spatial distribution of these effects was such that the emotional task was associated with higher MSE in finer timescales (Figure 5A) in bilateral bank of the superior temporal sulcus (bankSTS) and inferior parietal cortex; left hemisphere caudal and rostral middle frontal, and precentral regions; right mPFC, paracentral, pars triangularis, rostral ACC, precuneus, SP MT, and ST. The negative bootstrap ratios (Figure 5B) are reliable in bilateral insula, cingulate, ST, PCC, cuneus, and pericalcarine; left FG, mOFC, SF, parahippocampal, PCC, precuneus, and lingual; and right hemisphere MF, FP, OFC, IFG, postcentral, entorhinal, and temporal cortex.

**Figure 5.**
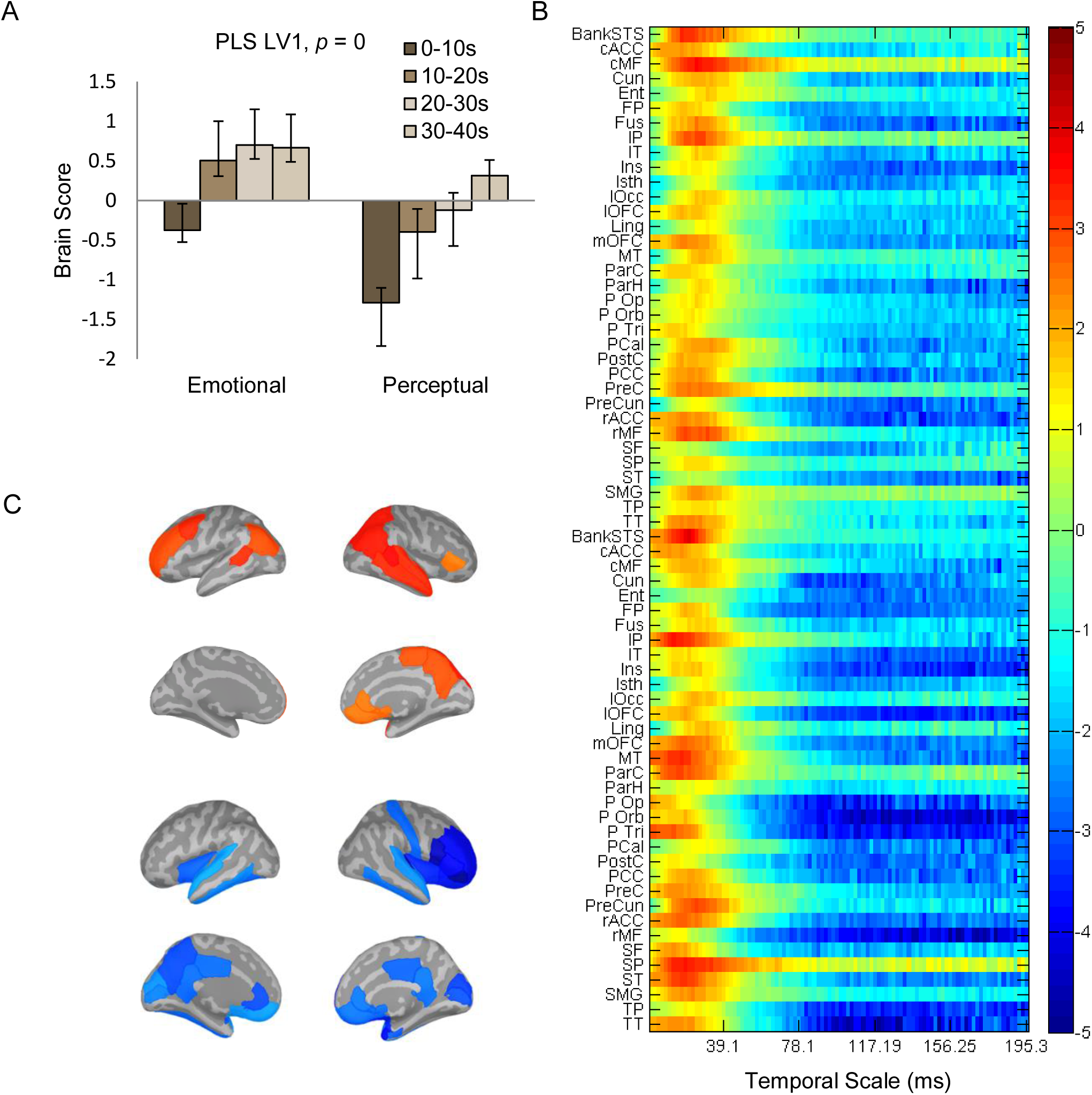
PLS first latent variable for the examination of effects of between tasks and within-task epochs on MSE. (A) The bar graph depicts the data-driven contrast highlighting higher MSE on all epochs of the emotional task compared to the perceptual task, as well as epoch effects within each task, significantly expressed across the entire data set, as determined by permutation tests. (B) Cortical regions at which the contrast was most stable as determined by bootstrapping. Values represent the ratio of the parameter estimate for the source divided by the bootstrap-derived standard error (roughly z scores). (C) Cortical visualization of stable bootstrap values for fine (top) and coarse (bottom) scales.

**Figure 6.**
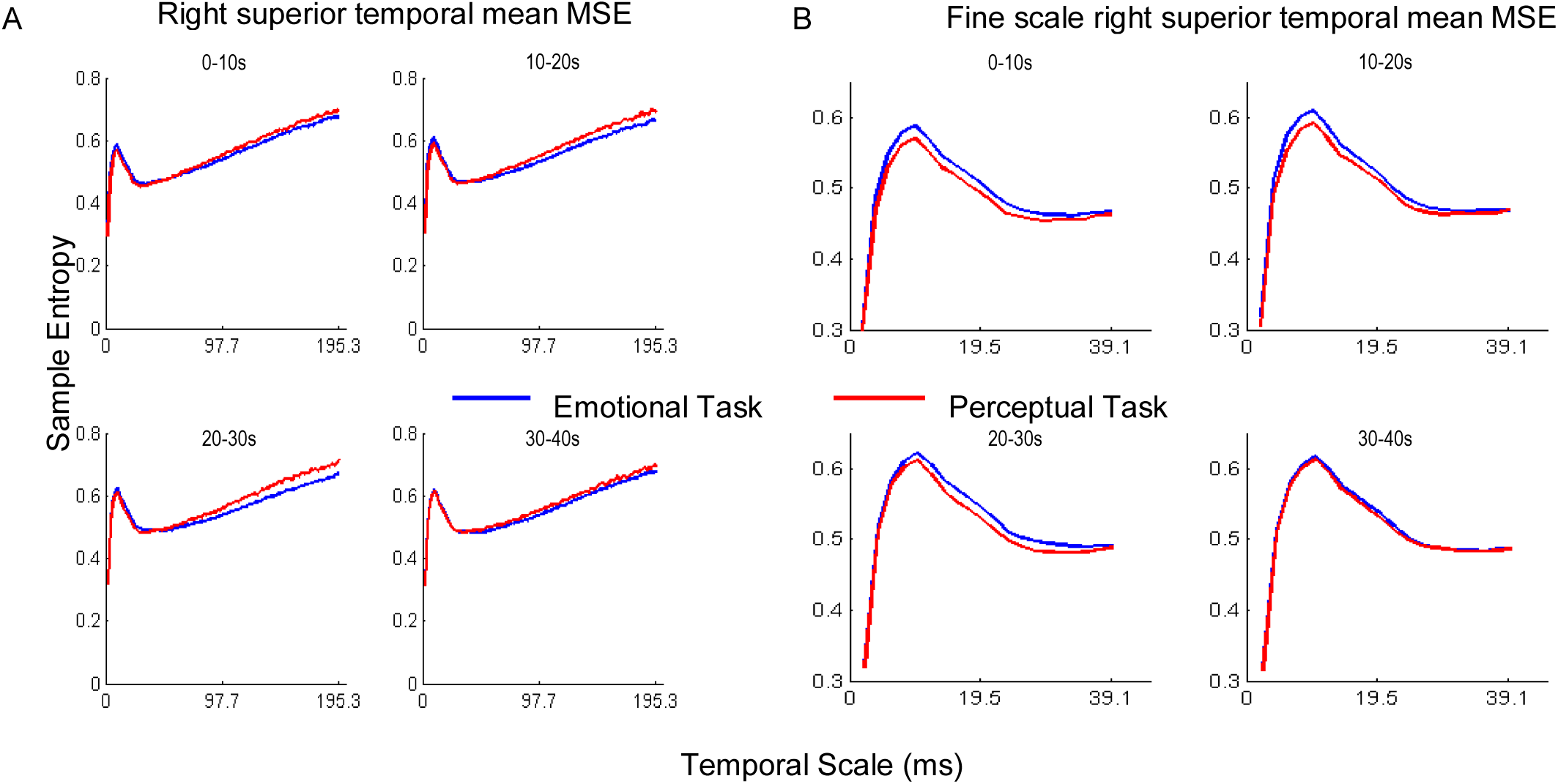
Mean MSE across participants for right superior temporal cortex ROI. (A) All temporal scales, B) Zoomed into visualize higher MSE for the emotional task at fine scales (<39.1 ms).

Procrustes’ distance was greater during the emotional task than the perceptual task in most brain sources (LV = 1, *p* = 0, Singular Value = 0.81, 54.1% of the cross-block covariance; Figure 7). All brain regions showed this effect of lower complexity matching on the emotional task except for bilateral entorhinal, FP, IT, parahippocampal, TP; left bankSTS MT, pars opercularis, ST; and right FG.

**Figure 7.**
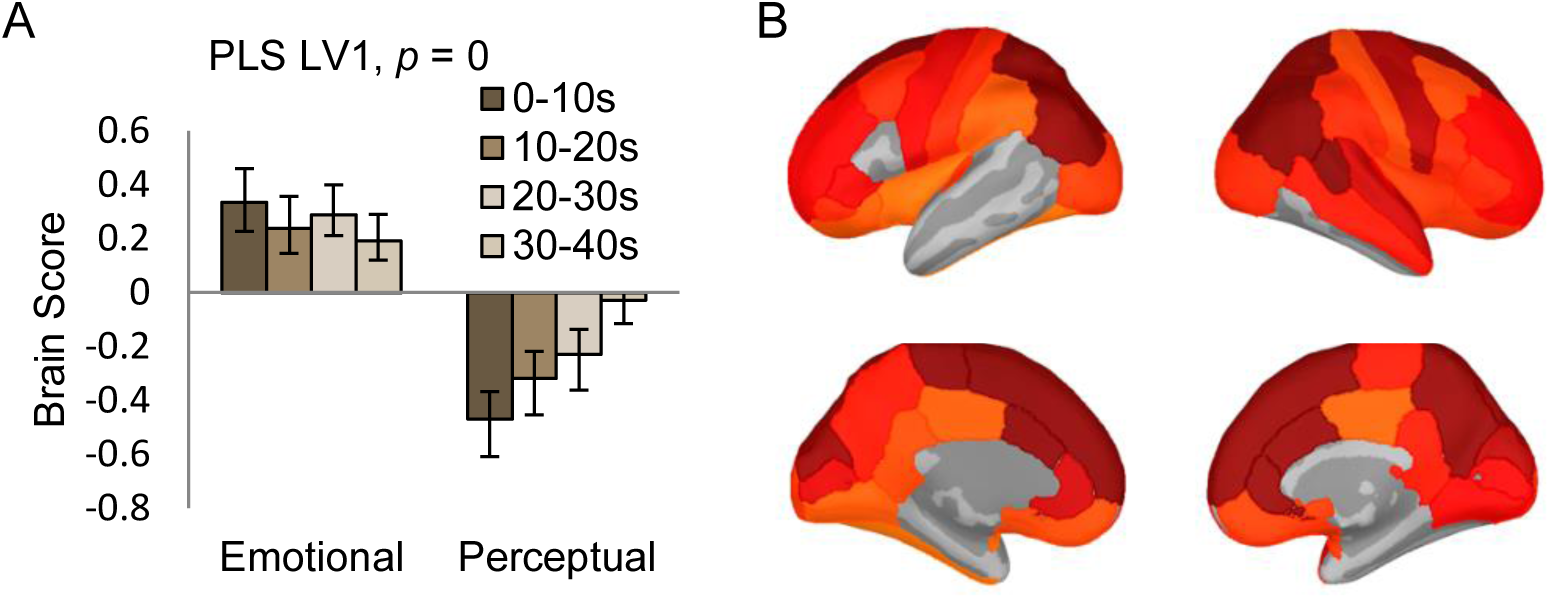
First significant PLS result for the examination of effects of between tasks and within-task epochs on Procrustes’ distance. (A) The bar graph depicts the data-driven contrast highlighting greater distance on all epochs of the emotional task compared to the perceptual task, as well epoch effects within each task, significantly expressed across the entire data set, as determined by permutation tests (p = 0). (B) Cortical regions at which the contrast was most stable as determined by bootstrapping. Values represent the ratio of the parameter estimate for the source divided by the bootstrap-derived standard error (roughly z-scores).

### Music Reward

The participant sample size (N=16) did not provide sufficient power to allow for the accurate assessment of brain-behaviour relationships on each of the five sub-factors of the BMRQ. Therefore, an average score across all sub-factors was calculated and used as the overall measure of music reward. Participant scores on this measure of reward had mean = 3.84 (SD = 0.47) on the 1-5 scale, suggesting this sample overall experiences a medium level of music related reward. Behavioural PLS assessed the correlation of the participant reward scores with MSE, distance and PSD on the two tasks and four epochs.

A strong positive correlation between MSE and reward was apparent during both tasks and all epochs. However, had we reported all epochs, the analysis would have included 8 conditions for only a total N=16; therefore we opted to not report the results of all epochs of both tasks in order to increase the validity of the statistical analysis and reduce the likelihood of a Type II error. Here we only report the positive correlation between MSE and reward during the first and last epoch of both tasks to demonstrate that the effect is relatively stable from the beginning to the end of the music (PLS LV1 *p* = 0, *r*^2^ = .38, Singular Value = 61.53, 74.4% of cross block covariance; Figure 8), and note that the pattern of effect similar for the middle two epochs. This effect was reliable in bilateral medial OFC, inferior frontal, cingulate, temporal and occipital regions, left precuneus and right superior frontal cortex.

**Figure 8.**
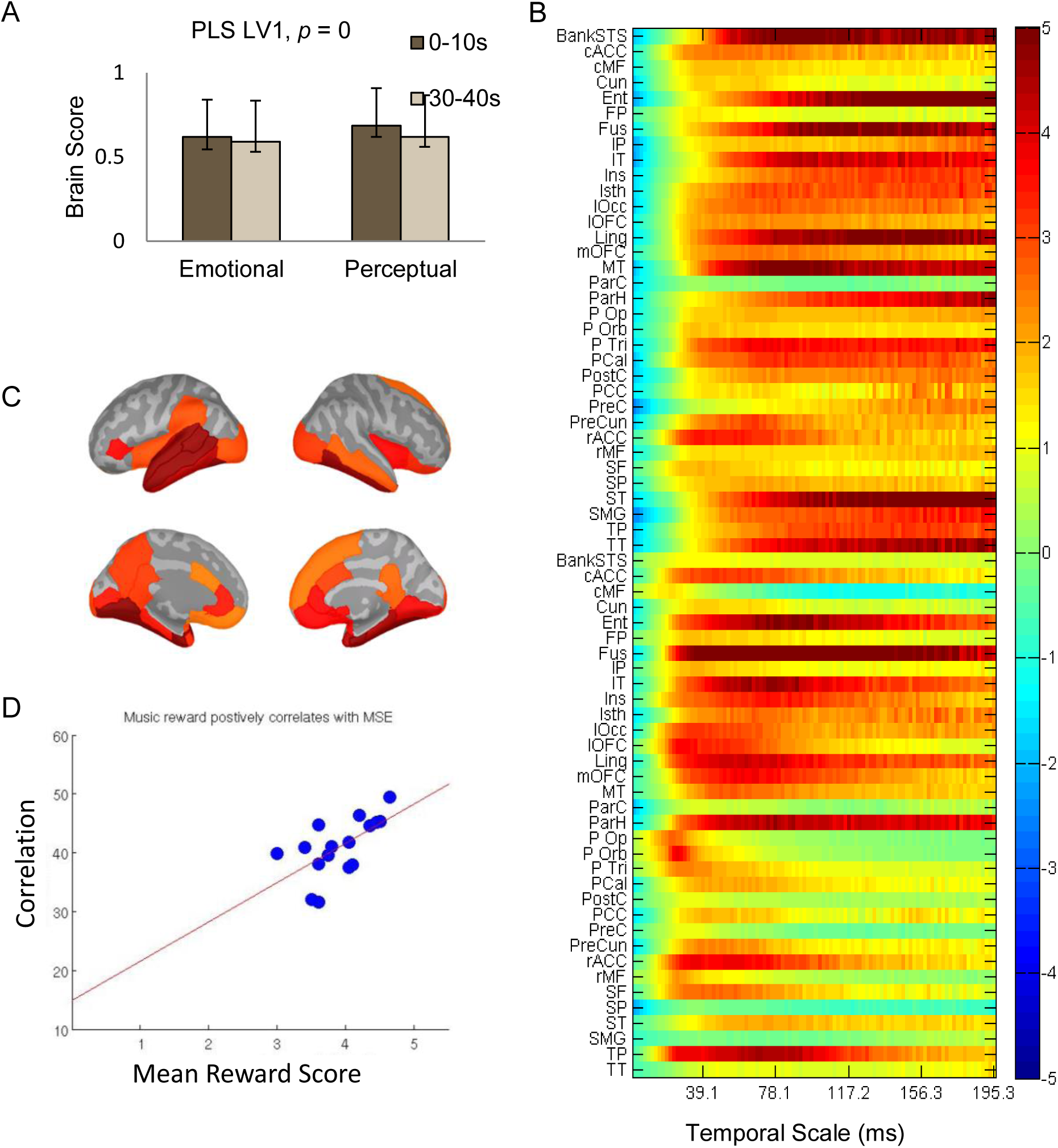
Behaviour PLS result examining the correlation between MSE and BMRQ reward score on both tasks and epochs E1 and E4. (A) Brain scores depict participants scores on the brain-behaviour relationship significantly expressed by the latent variable, as determined by permutation tests (p = 0). (B) Brain regions and frequencies at which the relationship was most stable as determined by bootstrapping. Together A and B indicate a positive correlation between MSE and BMRQ score in the highlighted regions. (C) Highlights bootstrap values from B for spatial regions where effect was stable. Values are taken as peak across scales 20-60. (D) Scatterplot of the brain scores from the first epoch with BMRQ reward scores depicts the positive relationship (*r*^2^ = .38).

A significant positive correlation was observed between distance and reward during only the emotional task for all epochs (PLS LV1 *p* = .012, *r*^2^ = .13, Singular Value = 4.29, 72.9% of cross-block covariance; Figure 9; perceptual task *p* > .10). This effect was localized to the right hemisphere frontal regions, rACC, IP, inferior and middle temporal, and lOcc.

**Figure 9.**
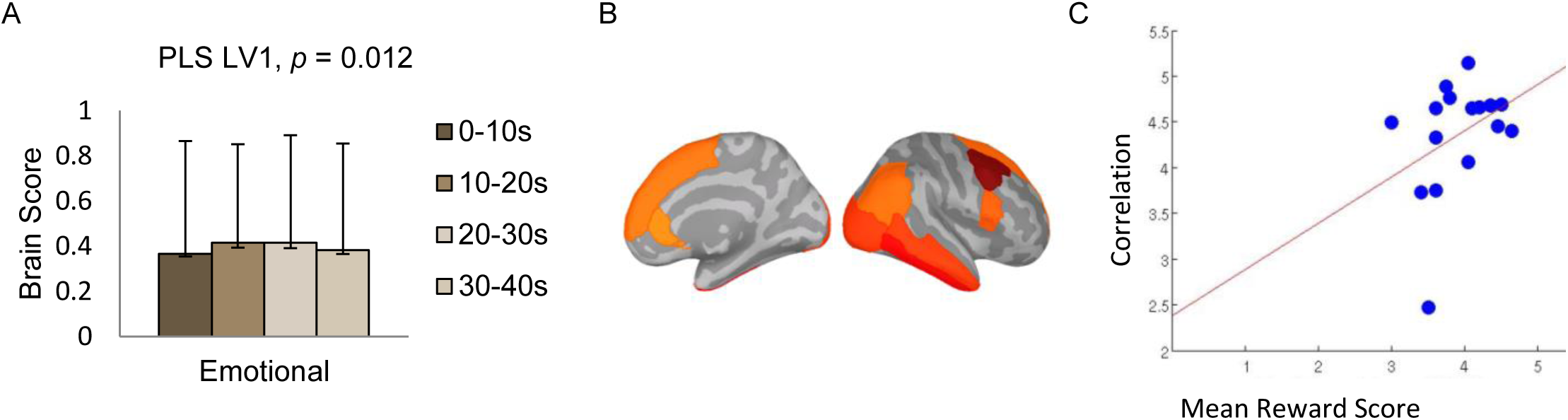
Behaviour PLS result examining the correlation between complexity distance on all epochs of the emotional task and BMRQ music reward score. (A) Brain scores depict participants score on the brain-behaviour relationship significantly expressed by the latent variable, as determined by permutation tests (p = .012). (B) Brain regions and frequencies at which the relationship was most stable as determined by bootstrapping. Together A and B indicate a positive correlation between distance and BMRQ score in the highlighted regions. (C) Scatterplot of the brain scores from the first epoch with BMRQ reward scores further displays this positive relationship (*r*^2^ = .13).

### Spectral Power

Higher gamma power was observed during the emotional task, and this effect increased across epochs (PLS LV1, *p* = 0, Singular Value = .40, 25.5% of cross-block covariance; Figure 10) in all spatial regions except for bilateral precentral gyrus and left pars triangularis. In comparison, the perceptual task was dominated by power at lower frequencies at the beginning of the piece of music, and this effect lessened over time.

**Figure 10.**
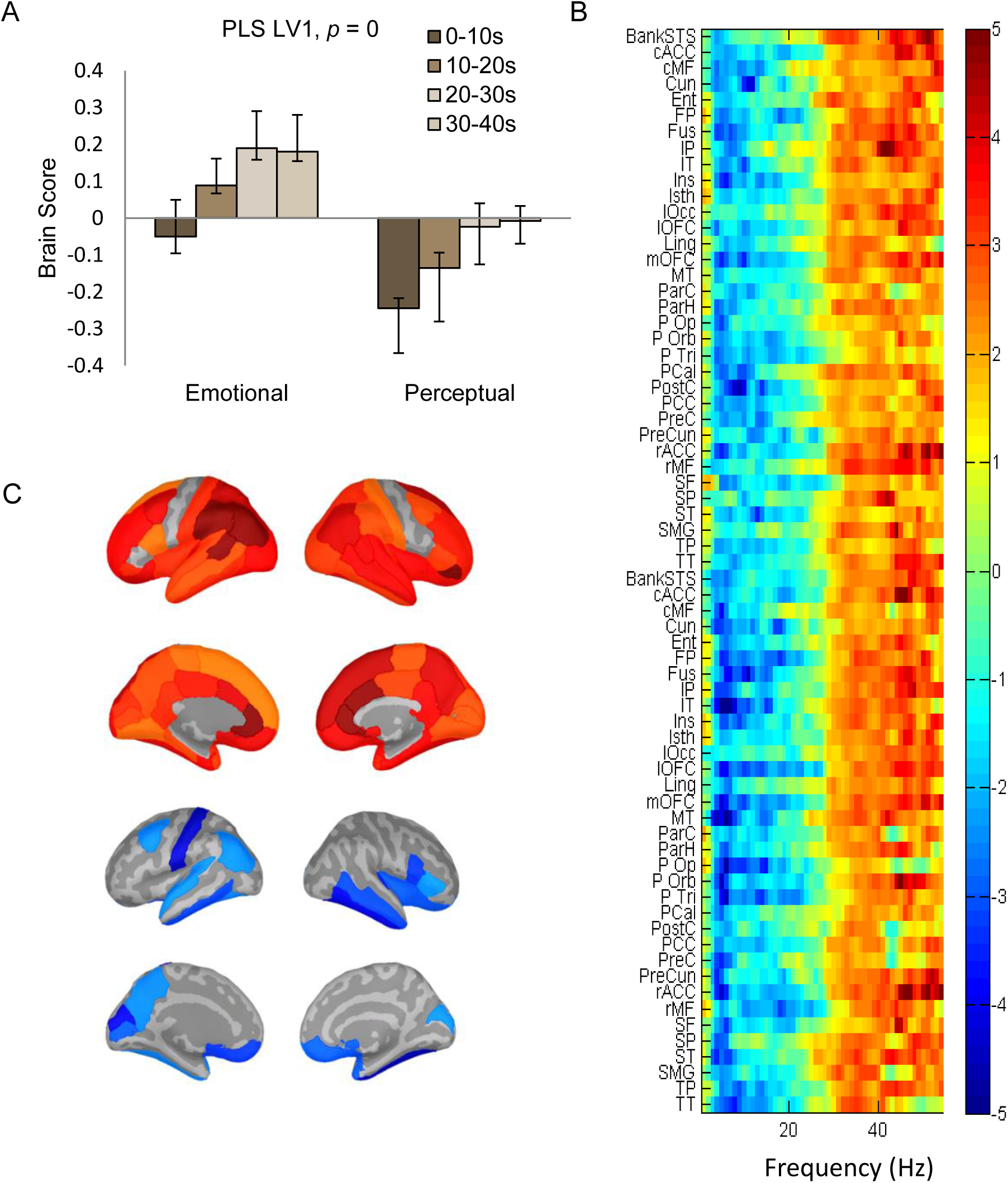
First significant PLS result for the examination of effects of between tasks and within-task epochs on PSD. (A) The bar graph depicts the data-driven contrast highlighting the differences in spectral power profile between tasks, and their similar epoch effect, significantly expressed across the entire data set, as determined by permutation tests (p = 0). (B) Cortical regions and frequencies at which the contrast was most stable as determined by bootstrapping. (C) Cortical visualization of stable bootstrap values (peak within each frequency band) for alpha (top) and gamma (bottom) band frequencies.

There was a positive correlation between music-derived reward as measured by BMRQ score and all spectral frequencies (PLS LV1, *p* = 0, *r*^2^ = .57, Singular Value = 44.49, 66.46 of cross-block covariance; Figure 11). The effect was spatially widespread across 58 of the 68 parcellated regions.

**Figure 11.**
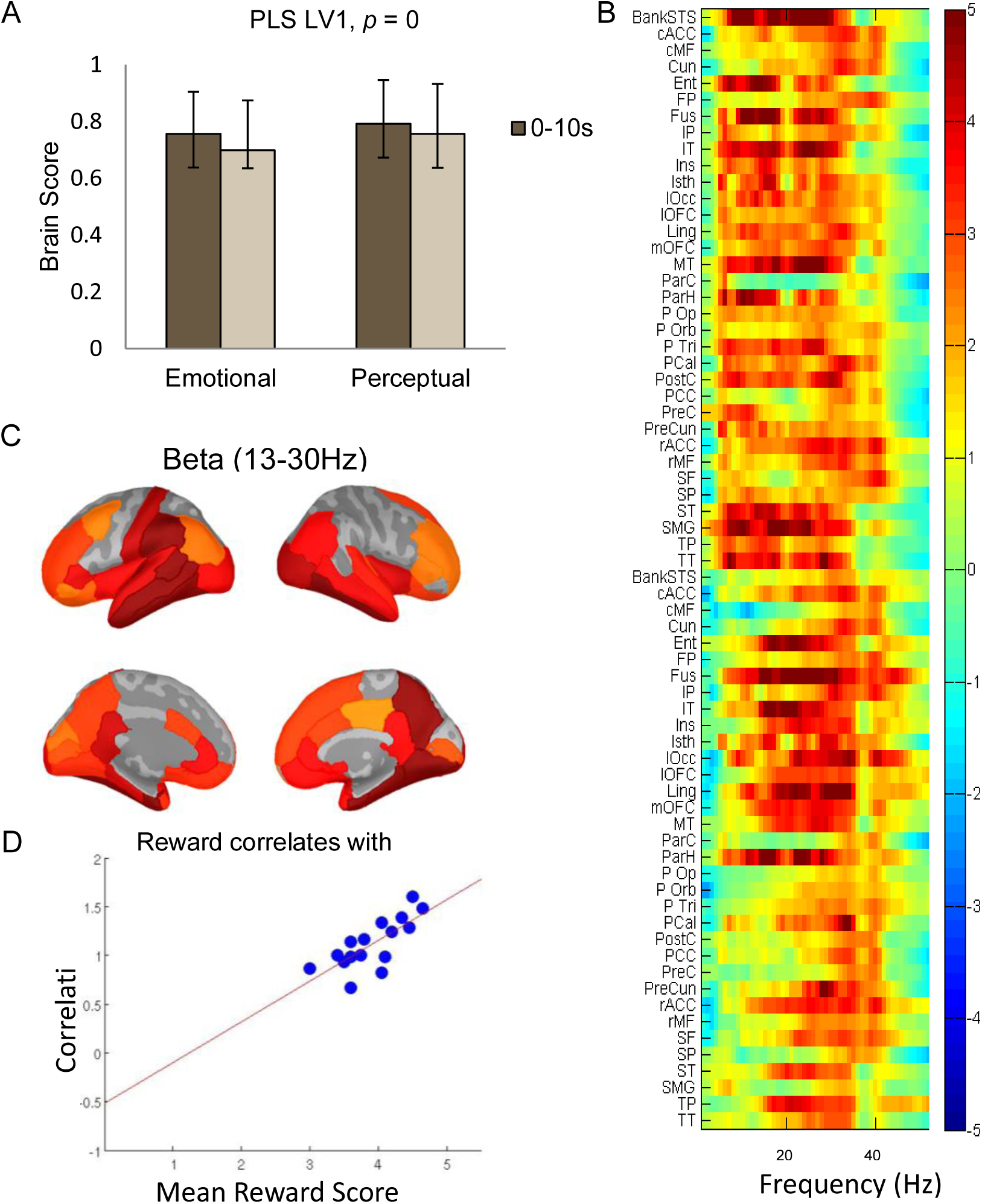
bPLS result examining the correlation between PSD and BMRQ reward score on both tasks and epochs E1 and E4. (A) Brain scores for the brain-behaviour correlation (p = 0). (B) Bootstrap ratios for brain regions and frequencies. Together A and B indicate a positive correlation between PSD and BMRQ scores across all frequencies in the highlighted regions. (C) Spatial regions from B where effect was stable in the beta band frequency (13-30 Hz). (D) Scatterplot of the brain scores from the first epoch with BMRQ reward scores depicts the positive relationship (*r*^2^ = .57).

## Discussion

We found higher complexity matching in widespread brain regions during the perceptual task than on the emotional task, using Procrustes’ distance to compare the MSE of EEG signals to the MSE of the music itself. This indicates that brain signal complexity more closely resembles the complexity of the music environment when participants were attending to the acoustics of the music compared to when they were thinking about how the music makes them feel. These results support a mapping of environmental information to the brain using complexity esatimation, and that the level of neural ‘mirroring’ is related to the type of cognitive processing conducted.

Analysis of the EEG MSE values alone found that MSE was higher in relatively finer timescales (<48.8ms) during the emotional task than the perceptual task, and that both groups showed an increase of this pattern as the music progressed. These MSE results demonstrate that emotion is associated with *higher information structure*, while the complexity matching results indicate the role of *different information structures* from environment input. The MSE results and the complexity matching results together suggest that the emotional task engaged additional processes, above and beyond the bottom-up sound perception information.

The brain regions that displayed this MSE effect are frequently linked to music cognition (e.g. right temporal, inferior frontal gyrus; Zatorre et al., 2002) and self-referential emotional processing (e.g. medial PFC; Amodio & Frith, 2006; Denny et al., 2012; Ochsner et al., 2004). The effect was also observed in regions that operate as integrative hubs (e.g. medial parietal) that are densely connected to neighboring regions and have long-range interconnections, enabling efficient global integration of information necessary for healthy cognitive function (Hagmann et al., 2008; Zamora-López et al., 2010; van den Heuvel and Sporns, 2011). This effect is spatially and temporally (<48.8ms) similar to increased complexity associated with musical training (Carpentier et al., 2016), and in other studies where higher MSE has been linked to performance on cognitive tasks that require higher information processing (Mišić et al., 2010; Heisz et al., 2012). Additionally, due to longer stimulus presentation, the current study opens up the analysis to coarser scales, up to 195.3ms, well beyond what has been possible in our previous studies and other empirical studies of cognition and brain MSE. We observed that MSE was initially higher in the perceptual task and decreased over time in the temporal scale range of 78.1-195.3ms, and we are not aware of any previous studies that have examined empirical brain complexity at this scale. We suggest the possibility that complexity at these slow timescales is indicative of an initial distributed and re-entrant search for neural templates associated with the music upon first listening, followed by a decrease of this activity as the brain settles into the neural solution for ongoing stimulus processing. However, further investigation of the link between cognition and complexity at these scales is required.

Higher music reward scores on the BMRQ were associated with a lower complexity match and higher EEG MSE. This negative correlation between complexity matching and reward was only observed on the emotional task, not on the perceptual task. Similar to the emotional task above, we suggest that reward is associated with additional internal information above and beyond that of the external stimulus. The complementary MSE and matching results suggest that music reward requires a brain state that is quantitatively and qualitatively different than the brain’s requirements for music sound perception alone. Music reward is associated with the activation of multiple different intrinsic processes and high information integration. In other words, music reward is a product of ‘the more you add’ to perception on top of immediate sensory events.

The observed relationship between higher neural information processing and music reward may be generated by the direct reward experience itself, since pleasurable responses to music are associated with particular patterns of cortical and subcortical activity not observed during neutral music perception. Multiple studies have reported connections between music reward and BOLD activity in vmPFC and OFC, and also IFG, ACC and sensory motor areas (Blood and Zatorre, 2001; Salimpoor et al., 2013). The relationship between complexity and reward in the present study was observed in temporal regions, as well as paralimbic and cortical regions involved in emotional processing (e.g. OFC, insula). Another proposal, not mutually exclusive to the first, is that the effects capture intermediate internal states that are important to generating the reward response. For example, the activity may reflect processes related to the BMRQ factors, such as musical knowledge, or other factors not directly measured by the BMRQ, like visual imagery or episodic memories evoked by the music (Juslin and Västfjäll, 2008; Vuoskoski and Eerola, 2012). Consistent with this notion, the spatial reliability of the correlation between distance and music reward suggest that frontal regions (superior, middle, inferior frontal) were processing internally generated information patterns, while inferior temporal and anterior cingulate cortex were involved in both the distance and MSE effects.

Integration may be a requirement for the commonly highlighted role of expectancy in music reward. The theory that rewarding emotional responses to music are derived from expectations and anticipation during music listening was first extensively described by Meyer (1956; see also Huron, 2006). It explains that the expectations are generated from explicit and implicit knowledge of music structure and patterns, and composers create emotional arousal by playing with ‘tension and release.’ Anticipation of a familiar rewarding segment of music has been linked to caudate dopamine release and BOLD activity prior to nucleus accumbens dopamine activity at peak reward response (Salimpoor et al., 2011). Music expectations were not behaviourally evaluated in the present study, but there is a logical link between them and brain signal complexity. Generation of expectations requires sufficient understanding and neural representation of the structure and patterns in the music. Therefore, it may be that enjoyment of a piece of music needs to be associated with a minimum amount of information processing that would allow the listener to appreciate the music as a coherent whole, rather as a sequence of individual notes. Further investigation of listeners’ enjoyment of individual music pieces, rather than as general trait music reward, is required to substantiate this theory and make a stronger connection between brain complexity and music pleasure.

While there were no notable differences in complexity between the song sets for each task, and the sets were selected to be acoustically and thematically similar, the songs were not identical for both tasks. This leaves open the possibility that other differences in the chosen songs are responsible for the observed brain differences between songs. This does raise interesting options for the future study of how different stimuli properties may influence brain complexity.

### Spectral Power

Congruent with the MSE pattern of the trade-off between faster and slower timescales, we observed higher gamma power in the emotional task, compared to lower frequency power associated with the perceptual task, as well as a decrease across time in low frequency power in regions typically linked to music processing and an increase in gamma in these and most other regions. Gamma activity has been repeatedly implicated as important for perceptual binding and may be associated with binding of musical features at the sensory level and matching of external acoustic information to internal thought processes for the formation of meaningful concepts (Bertrand & Tallon-Baudry, 2000; Crone et al., 2001; Keil et al., 1999; Rodriguez et al., 1999; Tallon-Baudry et al., 1998). Gamma activity is commonly found to be higher in adult musicians when listening to music and may reflect enhanced binding of musical features (Bhattacharya and Petsche, 2001, 2005; Shahin et al., 2008; Pallesen et al., 2015). There is also suggestion that gamma activity may be related to musical expectations (Snyder and Large, 2005). In a study conducted by Fujioka and colleagues (2009) gamma amplitude increased from baseline for each tone of a repeating pattern, and this effect continued on trials where the tone was unexpectedly omitted. None of these studies of the spectral effects of musical training or music listening conducted spatial analysis; therefore, it is difficult to place this facet of our results in the context of the other literature. However, our observation of increased gamma power in auditory and some associative regions is consistent with the hypothesis of the role of gamma in perceptual binding.

### Conclusions

EEG complexity was higher and different from music complexity during the emotional task in which participants were reflecting on how the music made them feel, compared to the perceptual task that had participants track pitch and tempo. Complexity matching was also correlated with BMRQ score, such that music reward was associated with higher neural signal information and a worse match to the bottom-up music information. These results suggest that complexity matching can assess the degree to which some cognitive-affective states are associated with internal information integration which differs from the neural representation of bottom-up sensory information processing.

## Acknowledgements

The authors wish to thank Natasa Kovačević for her work on the EEG preprocessing pipeline, as well as essential input at multiple levels of the Matlab scripting process.

## References

Amodio DM, Frith CD (2006) Meeting of minds: The medial frontal cortex and social cognition. Nat Rev Neurosci 7:268–277.

Bertrand O, Tallon-Baudry C (2000) Oscillatory gamma activity in humans?: a possible role for object representation. Int J Psychophysiol 38:211–223.

Bhattacharya J, Petsche H (2001) Musicians and the gamma band: a secret affair? Neuroreport 12:371–374.

Bhattacharya J, Petsche H (2005) Phase synchrony analysis of EEG during music perception reveals changes in functional connectivity due to musical expertise. Signal Processing 85:2161–2177 Available at: http://linkinghub.elsevier.com/retrieve/pii/S0165168405002070 [Accessed November 13, 2012].

Blood AJ, Zatorre RJ (2001) Intensely pleasurable responses to music correlate with activity in brain regions implicated in reward and emotion. PNAS 98:11818–11823.

Bressler SL, Kelso JAS (2001) Cortical coordination dynamics and cognition. Trends Cogn Sci 5:26–36 Available at: http://www.ncbi.nlm.nih.gov/pubmed/11164733.

Carpentier SM, Moreno S, McIntosh AR (2016) Short-term Music Training Enhances Complex, Distributed Neural Communication during Music and Linguistic Tasks. J Cogn Neurosci.

Carter FA, Wilson JS, Lawson RH, Bulik CM (1995) Mood Induction Procedure: Importance of Individualising Music. Behav Chang 12:159–161.

Catarino A, Churches O, Baron-Cohen S, Andrade A, Ring H (2011) Atypical EEG complexity in autism spectrum conditions: a multiscale entropy analysis. Clin Neurophysiol 122:2375–2383 Available at: http://www.ncbi.nlm.nih.gov/pubmed/21641861 [Accessed March 11, 2013].

Cohen AJ (2001) Music as a source of emotion in film. In: Music and Emotion: Theory and research (Juslin PN, Sloboda J, eds), pp 249–272. New York: Oxford University Press.

Costa M, Goldberger A, Peng C-K (2002) Multiscale Entropy Analysis of Complex Physiologic Time Series. Phys Rev Lett 89:6–9 Available at: http://link.aps.org/doi/10.1103/PhysRevLett.89.068102 [Accessed March 12, 2012].

Costa M, Goldberger A, Peng C-K (2005) Multiscale entropy analysis of biological signals. Phys Rev E 71:1–18 Available at: http://link.aps.org/doi/10.1103/PhysRevE.71.021906 [Accessed March 29, 2012].

Crone NE, Boatman D, Gordon B, Hao L (2001) Induced electrocorticographic gamma activity during auditory perception. 112.

Cross I, Morley I (2009) The evolution of music: theories, definitions and the nature of the evidence. In: Communicative musicality, (pp61–82) Oxford, Oxford University Press. (Malloch S, Trevarthen C, eds), pp 61–82. Oxford, UK: Oxford Univeristy Press. Available at: http://www.mus.cam.ac.uk/~ic108/PDF/CM_CM08.pdf.

Deco G, Jirsa VK, McIntosh AR (2011) Emerging concepts for the dynamical organization of resting-state activity in the brain. Nat Rev Neurosci 12:43–56 Available at: http://www.ncbi.nlm.nih.gov/pubmed/21170073 [Accessed October 26, 2012].

Delorme A, Makeig S (2004) EEGLAB: an open source toolbox for analysis of single-trial EEG dynamics including independent component analysis. J Neurosci Methods 134:9–21.

Denny BT, Kober H, Wager TD, Ochsner KN (2012) A Meta-analysis of Functional Neuroimaging Studies of Self- and Other Judgments Reveals a Spatial Gradient for Mentalizing in Medial Prefrontal Cortex. J Cogn Neurosci 24:1742–1752 Available at: http://www.mitpressjournals.org/doi/10.1162/jocn_a_00233.

Desikan RS, Ségonne F, Fischl B, Quinn BT, Dickerson BC, Blacker D, Buckner RL, Dale AM, Maguire RP, Hyman BT, Albert MS, Killiany RJ (2006) An automated labeling system for subdividing the human cerebral cortex on MRI scans into gyral based regions of interest. Neuroimage 31:968–980.

Efron B, Tibshirani R (1986) Bootstrap methods for standard errors, confidence intervals, and other measures of statistical accuracy. Stat Sci 1:54–77 Available at: http://www.jstor.org/stable/10.2307/2245500 [Accessed April 17, 2013].

Fujioka T, Trainor LJ, Large EW, Ross B (2009) Beta and gamma rhythms in human auditory cortex during musical beat processing. Ann N Y Acad Sci 1169:89–92.

Ghosh A, Rho Y, McIntosh AR, Kötter R, Jirsa VK (2008) Noise during rest enables the exploration of the brain’s dynamic repertoire. PLoS Comput Biol 4:e1000196 Available at: http://www.pubmedcentral.nih.gov/articlerender.fcgi?artid=2551736&tool=pmcentrez&rendertype=abstract [Accessed November 10, 2012].

Good P (2000) Permutation, Parametric and Bootstrap Tests of Hypotheses. Huntinton Beach, USA: Springer Science+Business Media, Inc.

Gower JC (1975) Generalized Procrustes Analysis. Psychometrika 40.

Gudmundsson S, Runarsson TP, Sigurdsson S, Eiriksdottir G, Johnsen K (2007) Reliability of quantitative EEG features. Clin Neurophysiol 118:2162–2171.

Hagmann P, Cammoun L, Gigandet X, Meuli R, Honey CJ, Wedeen VJ, Sporns O (2008) Mapping the structural core of human cerebral cortex. PLoS Biol 6:e159 Available at: http://www.pubmedcentral.nih.gov/articlerender.fcgi?artid=2443193&tool=pmcentrez&rendertype=abstract [Accessed November 3, 2012].

Heisz JJ, Shedden JM, McIntosh AR (2012) Relating brain signal variability to knowledge representation. Neuroimage 63:1384–1392 Available at: http://www.ncbi.nlm.nih.gov/pubmed/22906786 [Accessed March 11, 2013].

Holmes CJ, Hoge R, Collins DL, Woods R, Toda AW, Evans AC (1998) Enhancement of MR Images Using Registration for Signal Averaging. J Comput Assist Tomogr 22:324–333.

Hunter PG, Schellenberg EG (2010) Music Perception. In: Springer Handbook of Auditory Research (Jones MR, Fay RR, N PA, eds), pp 129–164. New York, USE: Springer.

Huron D (2006) Sweet Anticipation: music and the psychology of expectation. Cambridge, Massachusetts: MIT Press.

Juslin PN, Laukka P (2004) Expression, Perception, and Induction of Musical Emotions: A Review and a Questionnaire Study of Everyday Listening. J New Music Res 33:217–238 Available at: http://www.tandfonline.com/doi/abs/10.1080/0929821042000317813.

Juslin PN, Sloboda J (2010) Handbook of Music Emotions (Juslin PN, Sloboda J, eds). Oxford, New York: Oxford University Press.

Juslin PN, Västfjäll D (2008) Emotional responses to music: the need to consider underlying mechanisms. Behav Brain Sci 31:559–621 Available at: http://www.ncbi.nlm.nih.gov/sites/entrez?Db=pubmed&DbFrom=pubmed&Cmd=Link&LinkName=pubmed_pubmed&LinkReadableName=RelatedArticles&IdsFromResult=18826699&ordinalpos=3&itool=EntrezSystem2.PEntrez.Pubmed.Pubmed_ResultsPanel.Pubmed_RVDocSum.

Keil a, Müller MM, Ray WJ, Gruber T, Elbert T (1999) Human gamma band activity and perception of a gestalt. J Neurosci 19:7152–7161.

Krishnan A, Williams LJ, McIntosh AR, Abdi H (2011) Partial Least Squares (PLS) methods for neuroimaging: a tutorial and review. Neuroimage 56:455–475 Available at: http://www.ncbi.nlm.nih.gov/pubmed/20656037 [Accessed November 6, 2012].

Lippé S, Kovačević N, McIntosh AR (2009) Differential maturation of brain signal complexity in the human auditory and visual system. Front Hum Neurosci 3:48 Available at: http://www.pubmedcentral.nih.gov/articlerender.fcgi?artid=2783025&tool=pmcentrez&rendertype=abstract [Accessed April 30, 2013].

Mas-Herrero E, Marco-Pallares J, Lorenzo-Seva U, Zatorre RJ, Rodriguez-Fornells A (2013) Individual Differences in Music Reward Experience. Music Percept 31:118–138.

McIntosh AR (2000) Towards a network theory of cognition. Neural Networks 13:861–870 Available at: http://www.ncbi.nlm.nih.gov/pubmed/11156197.

McIntosh AR, Bookstein FL, Haxby J V, Grady CL (1996) Spatial pattern analysis of functional brain images using partial least squares. Neuroimage 3:143–157 Available at: http://www.ncbi.nlm.nih.gov/pubmed/9345485.

McIntosh AR, Kovačević N, Itier RJ (2008) Increased brain signal variability accompanies lower behavioral variability in development. PLoS Comput Biol 4:e1000106 Available at: http://www.pubmedcentral.nih.gov/articlerender.fcgi?artid=2429973&tool=pmcentrez&rendertype=abstract [Accessed November 20, 2012].

McIntosh AR, Lobaugh NJ (2004) Partial least squares analysis of neuroimaging data: applications and advances. Neuroimage 23 Suppl 1:S250–63 Available at: http://www.ncbi.nlm.nih.gov/pubmed/15501095 [Accessed November 5, 2012].

Meyer LB (1956) Emotion and Meaning in Music. Chicago, Il, USA: The University of Chicago Press.

Mišić B, Doesburg SM, Fatima Z, Videl J, Vakorin VA, Taylor MJ, McIntosh AR (2014) Coordinated Information Generation and Mental Flexibility: Large-Scale Network Disruption in Children with Autism. Cereb Cortex Available at: http://cercor.oxfordjournals.org/content/early/2014/04/25/cercor.bhu082%5Cnhttp://cercor.oxfordjournals.org/content/early/2014/04/25/cercor.bhu082.full.pdf%5Cnhttp://www.ncbi.nlm.nih.gov/pubmed/24770713%5Cnhttp://cercor.oxfordjournals.org/content/early/2014/04/.

Mišić B, Mills T, Taylor MJ, McIntosh AR (2010) Brain noise is task dependent and region specific. J Neurophysiol 104:2667–2676 Available at: http://www.ncbi.nlm.nih.gov/pubmed/20844116 [Accessed June 11, 2013].

Ochsner KN, Knierim K, Ludlow DH, Hanelin J, Ramachandran T, Glover G, Mackey SC (2004) Reflecting upon Feelings: An fMRI Study of Neural Systems Supporting the Attribution of Emotion to Self and Other. J Cogn Neurosci 16:1746–1772 Available at: http://www.mitpressjournals.org/doi/10.1162/0898929042947829.

Pallesen KJ, Bailey CJ, Brattico E, Gjedde A, Palva JM, Palva S (2015) Experience drives synchronization: The phase and amplitude dynamics of neural oscillations to musical chords are differentially modulated by musical expertise. PLoS One 10:1–21 Available at: http://dx.doi.org/10.1371/journal.pone.0134211.

Pascual-Marqui RD (2002) Standardized low resolution brain electromagnetic tomography (sLORETTA): technical details. Methods Find Exp Clin Pharmacol 24D:5–12.

Price CJ (2010) The anatomy of language: a review of 100 fMRI studies published in 2009. Ann N Y Acad Sci 1191:62–88 Available at: http://www.ncbi.nlm.nih.gov/pubmed/20392276 [Accessed September 16, 2013].

Richman JS, Moorman JR (2000) Physiological time-series analysis using approximate entropy and sample entropy. Am J Physiol - Hear Circ Physiol:H2039–H2049.

Rodriguez E, George N, Lachaux J-P, Martinerie J, Renault B, Varela FJ (1999) Perception’s shadow : long-distance synchronization of human brain activity. Nature 397:430–433.

Salimpoor VN, Benovoy M, Larcher K, Dagher A, Zatorre RJ (2011) Anatomically distinct dopamine release during anticipation and experience of peak emotion to music. Nat Neurosci 14:257–262 Available at: http://www.ncbi.nlm.nih.gov/pubmed/21217764 [Accessed May 22, 2013].

Salimpoor VN, van den Bosch I, Kovačević N, McIntosh AR, Dagher A, Zatorre RJ (2013) Interactions between the nucleus accumbens and auditory cortices predict music reward value. Science 340:216–219 Available at: http://www.ncbi.nlm.nih.gov/pubmed/23580531 [Accessed May 21, 2013].

Sampson PD, Streissguth AP, Barr HM, Bookstein FL (1989) Neurobehavioral effects of prenatal alcohol: Part II. Partial least squares analysis. Neurotoxicol Teratol 11:477–491 Available at: http://www.ncbi.nlm.nih.gov/pubmed/2593987.

Shahin AJ, Roberts LE, Chau W, Trainor LJ, Miller LM (2008) Music training leads to the development of timbre-specific gamma band activity. Neuroimage 41:113–122.

Snyder JS, Large EW (2005) Gamma-band activity reflects the metric structure of rhythmic tone sequences. Brain Res Cogn Brain Res 24:117–126 Available at: http://www.ncbi.nlm.nih.gov/pubmed/15922164 [Accessed January 30, 2013].

Tadel F, Baillet S, Mosher J, Pantazis D, Leahy RM (2011) Brainstorm: a user-friendly application for MEG/EEG analysis. Comput Intell Neurosci Available at: http://dl.acm.org/citation.cfm?id=1992539 [Accessed January 10, 2013].

Tallon-Baudry C, Bertrand O, Peronnet F, Pernier J (1998) Induced gamma-band activity during the delay of a visual short-term memory task in humans. J Neurosci 18:4244–4254 Available at: http://www.ncbi.nlm.nih.gov/pubmed/9592102.

Tononi G, Sporns O, Edelman GM (1994) A measure for brain complexity: relating functional segregation and integration in the nervous system. Proc Natl Acad Sci U S A 91:5033–5037 Available at: http://www.pubmedcentral.nih.gov/articlerender.fcgi?artid=43925&tool=pmcentrez&rendertype=abstract.

Tononi G, Sporns O, Edelman GM (1996) A complexity measure for selective matching of signals by the brain. Proc Natl Acad Sci U S A 93:3422–3427.

van den Heuvel MP, Sporns O (2011) Rich-club organization of the human connectome. J Neurosci 31:15775–15786 Available at: http://www.ncbi.nlm.nih.gov/pubmed/22049421 [Accessed May 21, 2013].

Västfjäll D (2001) Emotion induction through music: A review of the musical mood induction procedure. Music Sci 5:173–211 Available at: http://journals.sagepub.com/doi/10.1177/10298649020050S107.

Vuoskoski JK, Eerola T (2012) Can sad music really make you sad? indirect measures of affective states induced by music and autobiographical memories. Psychol Aesthetics, Creat Arts 6:204–213.

Zamora-López G, Zhou C, Kurths J (2010) Cortical hubs form a module for multisensory integration on top of the hierarchy of cortical networks. Front Neuroinform 4:1 Available at: http://www.pubmedcentral.nih.gov/articlerender.fcgi?artid=2859882&tool=pmcentrez&rendertype=abstract [Accessed June 13, 2013].

Zatorre RJ, Belin P, Penhune VB (2002) Structure and function of auditory cortex: music and speech. Trends Cogn Sci 6:37–46 Available at: http://www.ncbi.nlm.nih.gov/pubmed/11849614.

